# Structure of plant PSI-plastocyanin complex reveals strong hydrophobic interactions

**DOI:** 10.1101/2021.02.03.429574

**Authors:** Ido Caspy, Mariia Fadeeva, Sebastian Kuhlgert, Anna Borovikova-Sheinker, Daniel Klaiman, Gal Masrati, Friedel Drepper, Nir Ben-Tal, Michael Hippler, Nathan Nelson

## Abstract

Photosystem I is defined as plastocyanin-ferredoxin oxidoreductase. Taking advantage of genetic engineering, kinetic analyses and cryo-EM, our data provide novel mechanistic insights into binding and electron transfer between PSI and Pc. Structural data at 2.74 Å resolution reveals strong hydrophobic interactions in the plant PSI-Pc ternary complex, leading to exclusion of water molecules from PsaA-PsaB / Pc interface once the PSI-Pc complex forms. Upon oxidation of Pc, a slight tilt of bound oxidized Pc allows water molecules to accommodate the space between Pc and PSI to drive Pc dissociation. Such a scenario is consistent with the six times larger dissociation constant of oxidized as compared to reduced Pc and mechanistically explains how this molecular machine optimized electron transfer for fast turnover.

**One Sentence Summary:** Genetic engineering, kinetics and cryo-EM structural data reveal a mechanism in a major step of oxygenic photosynthesis

## Main Text

Biological energy conversion relies on a sequence of redox reactions within the membrane-embedded electron transport chains of chloroplasts and mitochondria. In plants, the photosynthetic electron transport between the two photosystems has to cover distances of a few hundred nanometers. This is due to unique architectural features of the photosynthetic thylakoid membranes, which fold into stacked grana and unstacked stroma thylakoids (*1*). However, the thylakoid architecture and the configuration of the protein complexes are subject to constant modulation in response to environmental and physiological stresses (*2-4*). Plastoquinone and plastocyanin (Pc) shuttles electrons from PSII to PSI, one within the hydrophobic milieu of the membrane and the second in the hydrophilic environment of the lumen (*5*). Pc exploits specific binding sites on its electron donor – cytochrome f - and its electron acceptor – PSI (*6*). The former interaction is ruder simple protein-protein interaction (*7*), and the latter is quite involved due to onset of P700 oxidation at the ns time scale and the electron transfer from Pc that take place at the μs time scale (*8-11*).

Binding and electron transfer between Pc and PSI is driven by hydrophobic and electrostatic interactions (*12*). In plants and green algae, the electron transfer reaction between Pc and PSI exhibits two distinct kinetic phases. A first fast microsecond intra-molecular electron transfer can be explained by a stable complex between PSI and Pc already formed prior to a flash. In the second slower phase, the remaining PSI complexes are re-reduced by the soluble donor in a bimolecular reaction with second-order kinetics. The occurrence of a pre-formed complex is dependent on the presence of the eukaryotic positively charged domain of subunit PsaF. The dissociation constant of oxidized Pc is about six times larger than that observed for reduced Pc (*13*), resulting in a 50-60 mV higher midpoint redox potential of Pc bound to PSI as compared to soluble Pc. This translates into a decrease of the driving force within the intra-molecular electron transfer complex, which is consistent with the fact that the P700 population is not fully reduced in vivo (*14*). Moreover, single turnover flashes of isolated PSI fail to oxidize the whole P700 population (*15, 16*).

In this work we used the single particle cryo-EM technique to obtain a high-resolution structure of reduced Pc in complex with plant PSI and analyzed the interaction between the two complexes. For kinetic studies, a highly purified PSI derived from PSI crystals and purified Pc were employed.

## Results

### Structure of reduced Pc in complex with PSI

The availability of large amounts of *Pisum sativum* PSI crystals that are stable for years in the crystalline form have offered a unique opportunity to solve the structure of photosynthetic supercomplexes by the cryo-EM technique (*17*). We have utilized these crystals to form a plant Pc-PSI complex simply by adding purified Pc to pure PSI. To ensure the complex formation the concentration of Pc was kept way above the dissociation constant. Plant PSI was solubilized from stored crystals in a solution containing 20 mM MES-Tris (pH7), 20 mM sodium ascorbate and 0.05% αDM at a chlorophyll concentration of 6 mg/ml. Concentrated pea Pc was added into the PSI solution to give final concentration of 2.8 mg Chl/ml PSI and 5 mg/ml Pc. The resulting solution of PSI-Pc (3 µl) was applied on glow-discharged holey carbon grids that were vitrified for cryo-EM structural determination (see Methods). Following data processing (see Methods section) the PSI-Pc supercomplex was solved at 2.74 Å resolution with local resolution ranging from 2.5 to 4.5 (fig S1, table S1, PDB 6ZOO). The structure of Pc-PSI complex is shown in Figure 1. The position on PSI of the reduced Pc is virtually identical to its position in the previously reported Pc-PSI-Fd complex (*17*; PDB 6YEZ). However, improved map densities of Pc allowed a closer look not only at the electrostatic but also potential hydrophobic interactions (Fig 1; movie S1; PDB 6ZOO). Figure 1 shows some of the identified surface interactions where previous molecular biology studies revealed deleterious mutations, with PsaA Trp651 and PsaB Trp627 (Trp658 and Trp625 in pea, respectively) at the center of the hydrophobic binding region (*18, 19*). Mutagenizing PsaB Glu613 and Asp612 (pea Glu611 and Asn612), which are located in the periphery of Pc binding domain, increased the affinity of Pc binding to PSI alongside with slower Pc release (10, *28*). PsaF lysines were observed in close proximity to Pc negatively charged amino acids as reported in (*17*), with the addition of PsaA Arg117 identified 5.7 Å away from Pc Asp61. Substitution of these lysines disrupted Pc binding, release, and electron transfer to P700 (*20*). On the Pc hydrophobic surface, mutating Pro36 - which faces the two tryptophans - as well as Gly10, Leu12, Tyr83 and Ala90 (*8, 22, 23*) showed a decreased efficiency in Pc binding and activity. The hydrophobic Pc region involved in PSI-Pc association was suggested to serve a similar role in Pc binding to cytochrome f (*24, 25*).

**Fig 1.**
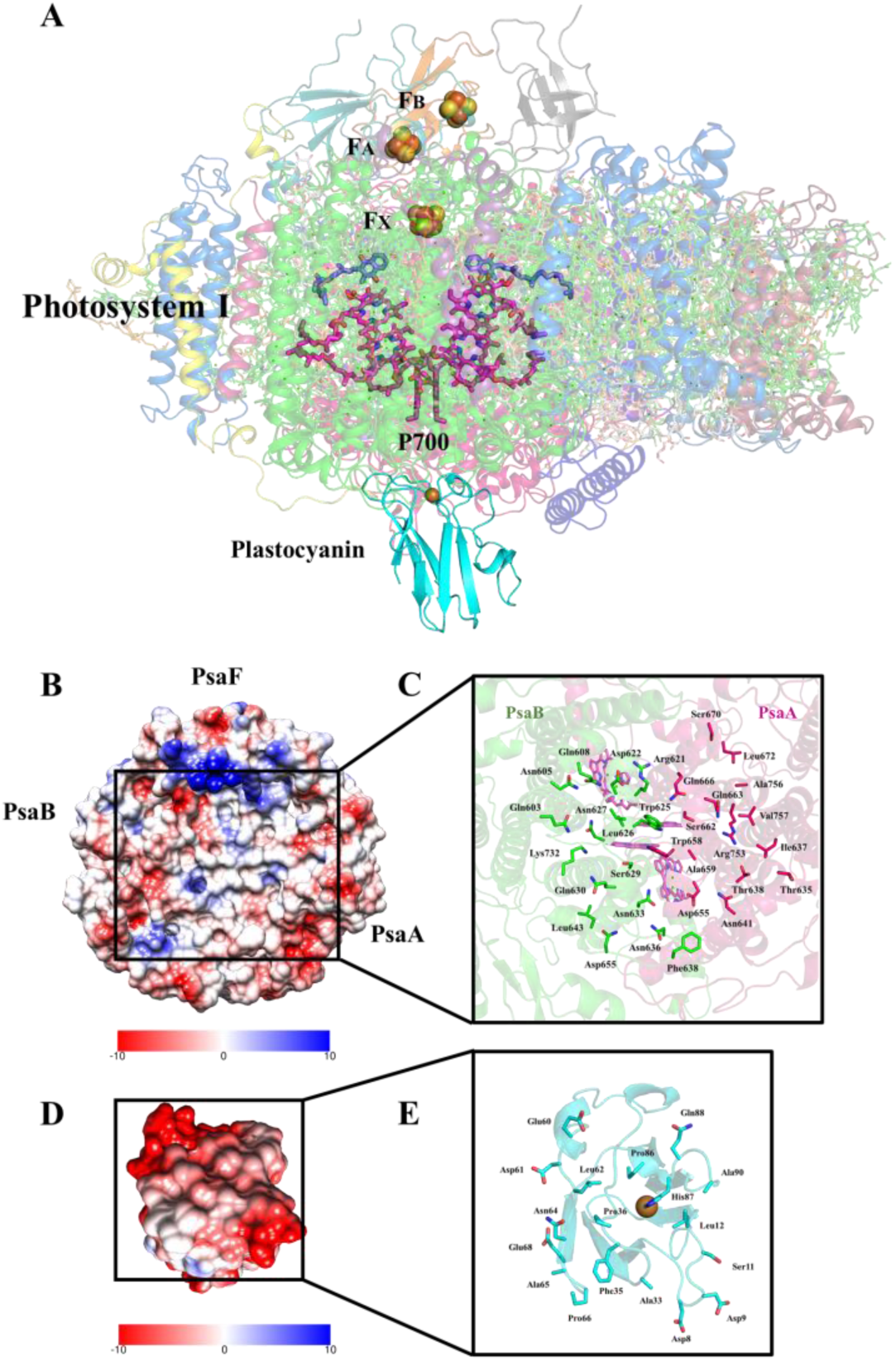
Structure of Photosystem I in-complex with reduced Pc and analysis of the solvent inaccessible region formed by their bound interfaces. (A) Side view of PSI-Pc complex. (B) Electrostatic potential analysis of the Pc binding surface formed by PsaA-PsaB. (C) Zoom-in on the PsaA-PsaB amino acids forming the hydrophobic Pc binding surface. (D) Electrostatic potential analysis of the Pc region that binds PSI. (E) Zoom-in of the Pc amino acids that make up the water inaccessible surface across PsaA-PsaB upon PSI-Pc complex formation.

### Binding surfaces of PSI-Pc and in-silico modelling of Pc release from PSI

While structural information readily reveals electrostatic interactions, hydrophobic interactions are much more difficult to decipher (*17*). One of the hallmarks of such interaction are the close contacts of the hydrophobic surfaces leaving no space for water molecules. Analysis of the solvent-accessible surface area (SASA) of the Pc binding domain formed by PsaA-PsaB and Pc surface using GetArea with default parameters (*26, 27*) demonstrates that more than 50% of the binding regions become inaccessible to water molecules once the PSI-Pc complex forms, fitting a tightly packed hydrophobic binding (Fig 1, B to E, Table 1). These regions are composed of four domains in Pc and five domains in PsaA and PsaB are presented in Figure 1, and several of these amino acids were previously identified as necessary for the transient binding of Pc (*8, 9, 17, 18, 19, 28*). These binding domains display a combination of electrostatic, polar, and hydrophobic interactions with varying magnitudes. Pc domains span from Asp8-Leu12, Ala33-Pro36, Glu60-Leu62, Asn64-Glu68 and Pro86-Gly91. PsaA and PsaB association domains span more intermittently but form five distinct surfaces encircling PsaA Trp658 and PsaB Trp625. The first PsaA surface is formed by Thr635, Ile637-Gly639 and Asn641, the second by Asp655, Trp658, Ala659, Ser662, Gln663, Gln666, Gly669, Ser670 and Leu672 and the third by Arg753, Ala756 and Val757. PsaB first domain consists of Gln603, Asn605 and Gln608, and the second by Arg621, Asp622, Trp625-Asn627, Ser629, Gln630, Asn633, Asn636, Phe638, Asn641, Leu643 and Lys732.

**Table 1.**
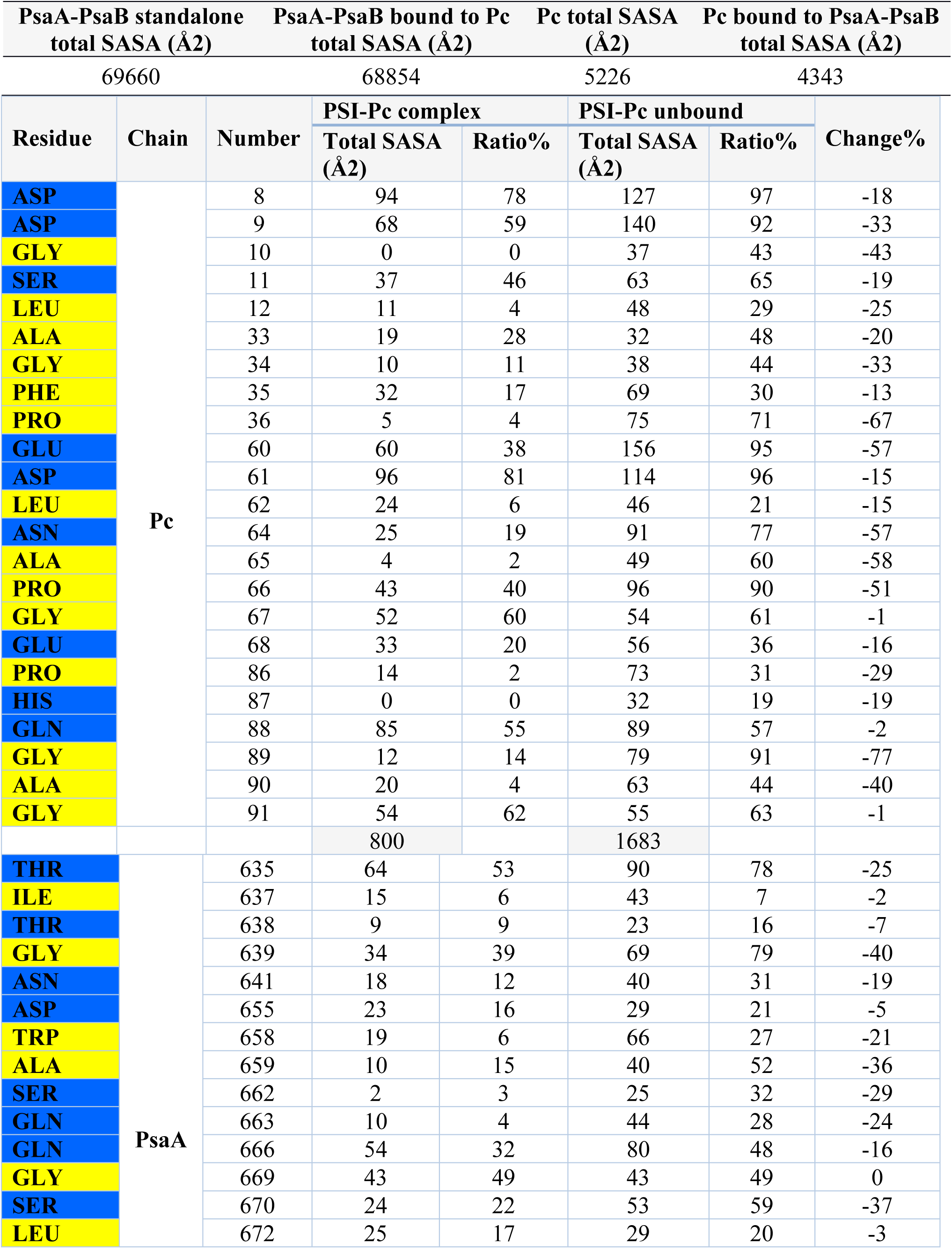
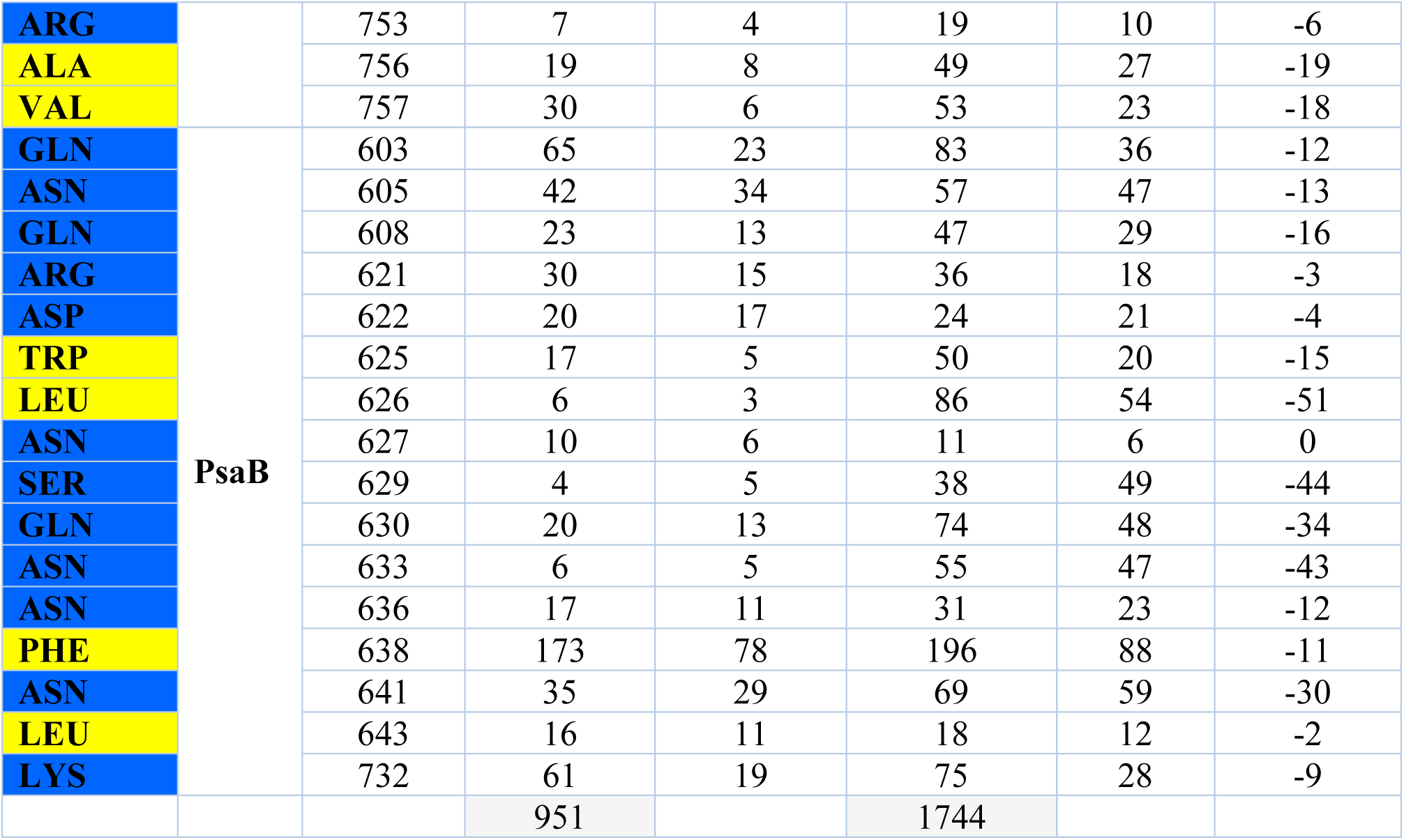
Solvent accessible surface area (SASA) analysis of Pc, PsaA and PsaB showing a hydrophobic binding area formation during PSI-Pc association. Total - SASA for each residue in Å^2^; Ratio – percentage of the area of residue exposed to the solvent; Change – difference in SASA for each residue, in complex or unbound. Charged or polar amino acids are colored blue, hydrophobic amino acids colored yellow.

As part of our efforts to describe Pc association and dissociation mechanism, we used in-silico modelling of oxidized Pc from *Populus nigra* (*29*; PDB 4DP9) as our reference. Sequence comparison of Pc from *P. nigra* and pea using BLAST (*30*) shows a 79% identity (fig S2). The sequence was changed according to pea Pc and then superpositioned on the reduced Pc in 6ZOO. Conformation changes which appeared significant were identified in both Pc acidic patches Asp42-Glu45 and Glu59-Glu60 (Fig 2). The short helix formed by Asp42-Glu45 in oxidized Pc brings Glu43 and Asp44 closer to PsaF Lys93, Lys96 and Lys100, and Glu60 drifts closer to PsaF Lys101. Next, we superimposed the two conformations based on unchanged core region (Asp61-Asn64). In this superposition the negative patches mentioned before were closer to PsaF Lysines, and tilted relative to the hydrophobic binding region. With this, Pro86-Gly91 moved by 1-4 Å compared to the 6ZOO conformation, leaving ample room for water molecules to accommodate the space between Pc and PSI and drive Pc dissociation from PsaA-PsaB towards PsaF (Fig 2, movie S2). Inspection of the in-silico model SASA shows that the oxidized Pc surface solvent accessibility increased by 5% compared to the reduced Pc, mostly in the hydrophobic binding patches (table S2).

**Fig 2.**
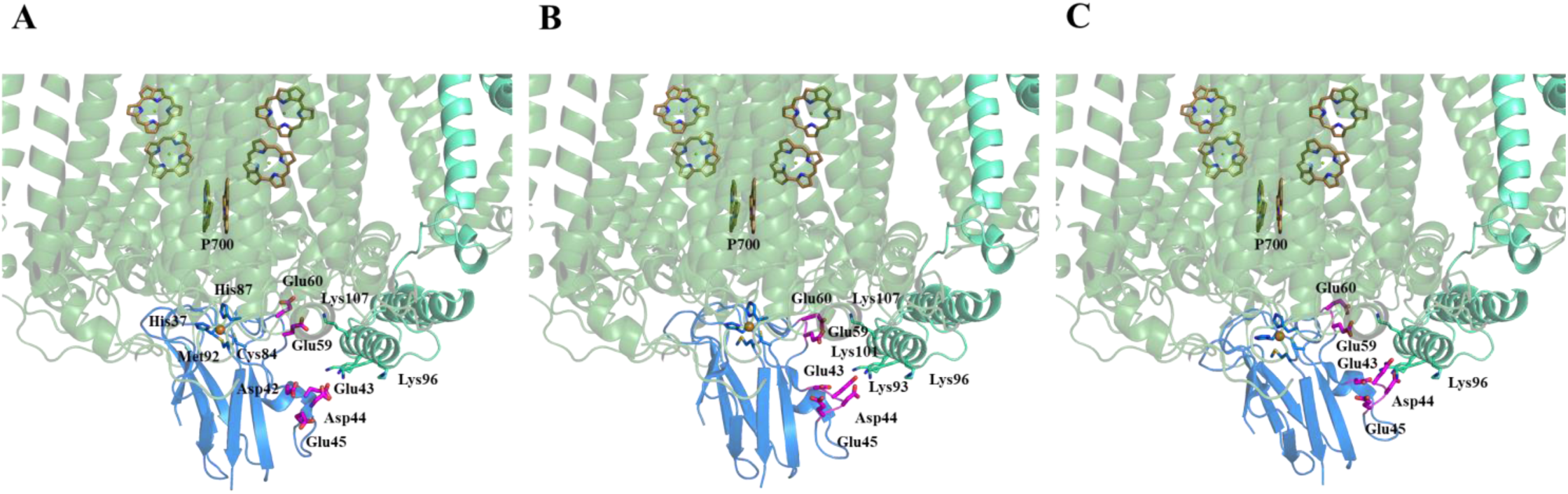
In-silico model of Pc dissociation from PSI. (A) Reduced Pc bound to PSI. Pc acidic patch is colored in magenta, Pc is colored marine, PSI in forest, PsaF in greencyan and P700 in sand. (B) Oxidized Pc (PDB 4DP9) bound to PSI in-silico. Acidic patches undergo a conformational change, bringing them closer to PsaF positive Lysines. (C) Superposition of unchanged regions Asp61-Asn64 causes Pc to tilt and permits water molecules to enter the space between the proteins.

### PsaA residues R647 and D648 are required for efficient binding and electron transfer between PSI and Pc

To further assess the role of amino acids close to the hydrophobic core represented by PsaA-Trp658, residues PsaA-Arg647 and Asp648 (Arg654 and Asp655 in pea, respectively) were altered via a site-directed mutagenesis approach using *Chlamydomonas reinhardtii* as a model. PsaA-Asp648 faces Pc Phe35 directly (fig S3, see supplementary text). SDS-PAGE fractionation of isolated thylakoid membranes followed by an immuno-blot analysis using anti-PsaD and anti-PsaF antibodies revealed diminished PSI contents of about 25 % to 75 % as compared to wild type, while the PSII content was stable (fig S4).

The second order electron transfer rates for the electron transfer between Pc and purified PSI complexes were determined via *in vitro* single turn-over flashes for excitation at increasing ionic strength (Fig 3A). All genetically modified photosystems revealed a diminished second order transfer rate as compared to wild type, independent of the salt concentration, with the strongest impact for PsaA-Asp648Arg (table S3).

**Fig 3.**
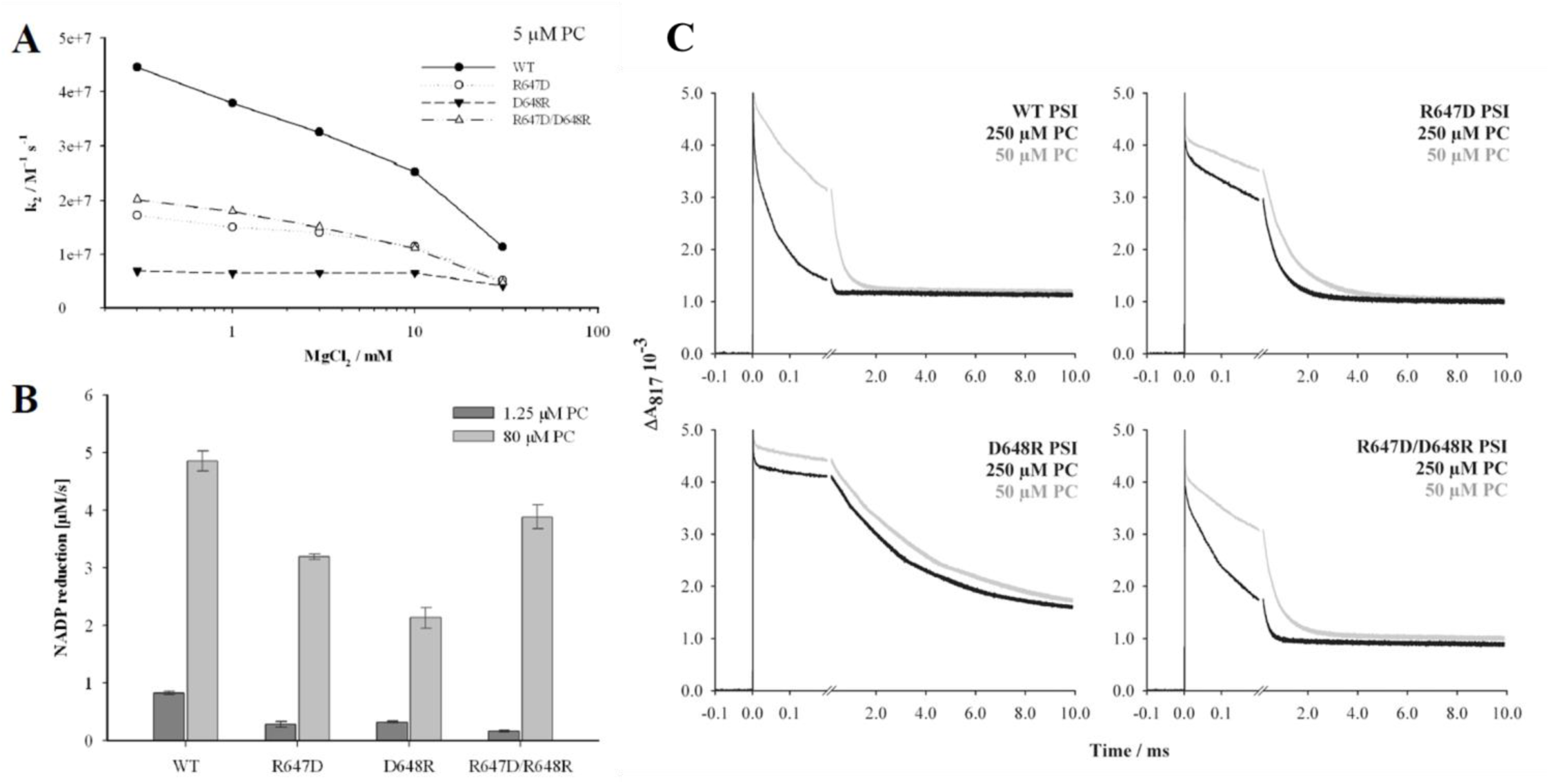
PsaA residues Arg647 and Asp648 are required for an efficient binding of the electron donor Pc. (A) Second-order rate constant of P700+ reduction by pc (5 μM) at increasing MgCl2 concentrations. The ionic strength is increased by adding small amounts of concentrated MgCl2 solution. The optimal salt concentration is at 1 mM MgCl2 for wild type and altered PSI, respectively. The second order electron transfer rate is decreased for all altered photosystems in comparison to wild type. (B) Light driven PSI dependent NADP+ photoreduction measurements for wild type and altered photosystems with pc at low (1.25 μM) and high concentration (80 μM). (C) Absorbance transient at 817 nm of the flash-induced photooxidation of wild type and the altered photosystems R647D, D648R and R647D/D648R and their subsequent re-reduction in the presence of 50 and 250 μM pc. The first order phase shows a half time about 4 μs, thus the relative amplitude (A1) is smaller in all altered photosystems as compared to wild type (increase of A1/(A1+A2) from 50 to 250 μM pc, for WT 100%, for R647D 45%, for D648R 17% and for R647D/D648R 30 %).

These findings were supported by light-induced PSI dependent NADP^+^ reduction measurements in an independent approach. At low Pc concentrations (1.5 µM) the photo-reduction rate was about 2-3 times (PsaA-Arg647Asp, PsaA-Asp648Arg) up to 5 times smaller (PsaA-Arg647Asp/Asp648Arg) for altered PSI was lower as compared to wild type (Fig 3B).

Similar effects were observed at high Pc concentrations (80 µM), yet the difference was only about 1.5 times smaller for PsaA-Arg647Asp and PsaA-Arg647Asp/Asp648Arg and 2.5 times smaller for PsaA-Asp648Arg. Both independent approaches indicated an impaired electron transfer from Pc to the altered PSI, independent from the residue combination.

Absorbance transients at 817 nm, induced by fast single turn-over flashes, were also recorded at increasing Pc concentrations (Fig 3C). The absorbance transients at high Pc concentrations can be separated into three kinetic components. The fast component A(1) with a constant half-life of 3-4 µs representing the first order electron transfer reaction within a preformed complex, a second components A(2) with a half-life decreasing at higher concentrations indicative of a second-order binding reaction, and a very slow third component A(3) representing PSI complexes that lost their PsaF subunit during the isolation process making up to 25-40 % of the total PSI signal (*31*). The changes in amplitude of the two faster kinetic components in regard to the donor concentration reflect the binding equilibrium and allows the calculation of the dissociation constant K_D_ according to Drepper et al. (*13*). The K_D_ value for WT PSI was measured at 88 μM, similar to previous work (*20*). Determination of the K_D_ values for all altered photosystems was not possible as the amplitude A(1) did not significantly increase at increasing Pc concentrations, visible in the absorbance transients measured in the presence of 50 and 250 µM Pc (Fig 3C). This reflects a drastically decreased affinity of Pc for the altered photosystems, indicating that changing charges at PsaA residues Arg647 and Asp648 omitted efficient pre-binding of reduced Pc.

### Reduction kinetics of light induced P700+

Electron transfer measurements were performed using a uniform PSI population derived from solubilized diffracting crystals to substantiate that these PSI complexes are very competent in electron transfer. The light-induced P700 oxidation and re-reduction by various electron donors were followed by recording absorption changes at 705 nm using a Joliot-Type Spectrophotometer (JTS). The signals converted to assume a mM extinction coefficient of 64 at 705 nm (*32*). No significant change in OD was observed during illumination from 0.1 to 1 sec (fig S5). Similar to ferredoxin and PMS, the presence of methylviologen also eliminated the fast kinetics (Fig 4A). Addition of NaCl significantly decreased the rate of P700+ reduction by Pc (Fig 3A, Fig 4B). Pre-incubation with sufficient plastocyanin amounts should saturate the plastocyanin binding site even under conditions where the electrostatic interactions are eliminated by the presence of high salt concentration. Measurements at a shorter time scale revealed that the A(1) phase, the initial P700+ reduction by pre-bound Pc, is insensitive to the presence of high salt concentration in the reaction mixture (Fig 4C). Under these conditions hydrophobic interactions dominate the Pc binding to PSI, underpinning the importance of such interactions between PSI and Pc.

**Fig 4.**
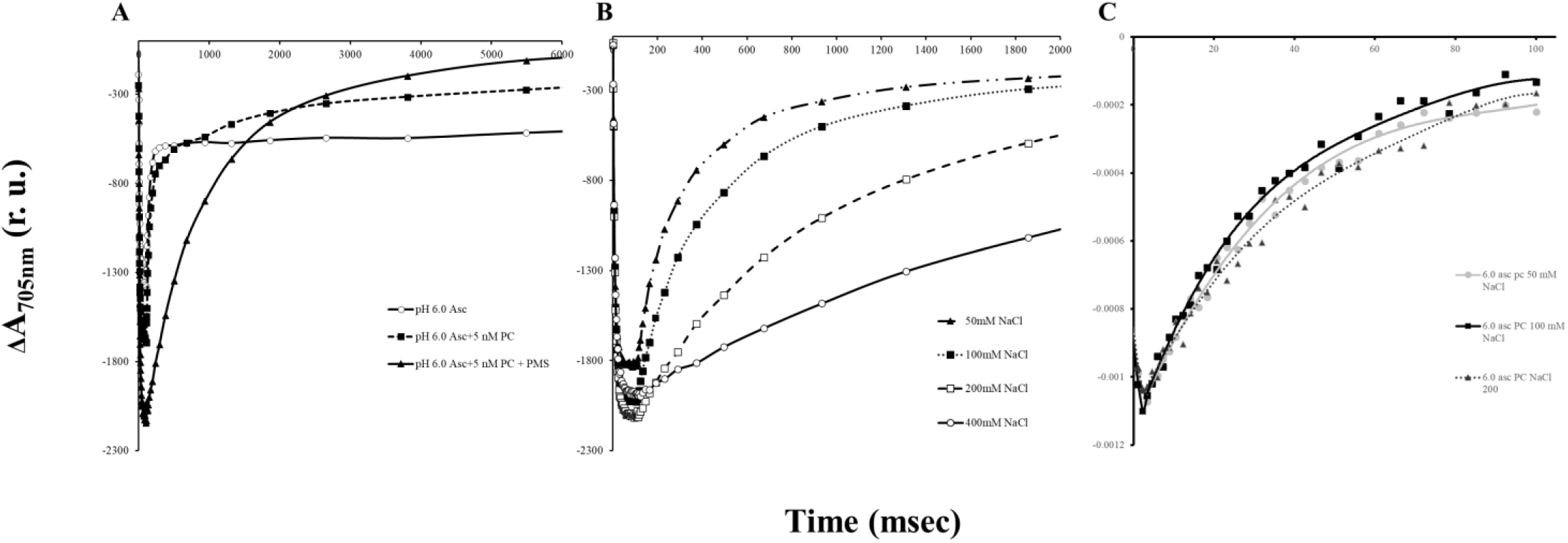
P700 reduction by Pc is dependent on ionic strength and solvent pH. (A) Kinetics of P700+ reduction in plant PSI derived from dissolved crystals in the presence of ascorbate and PMS. Reaction mixture contains: 20 mM MES-Tris (pH=6.0); 0.05% α-DM and PSI containing 5 µg Chl. Ascorbate – 2 mM; Pc - 5 nM; PMS - 1 μM. Excitation light for 0.1 sec. (B) Effect of NaCl concentration on the rate of P700^+^ reduction by Pc. Reaction mixture contains: 20 mM MES-Tris (pH=6.0); 0.05% α-DM and PSI containing 5 µg Chl. All samples contain 10 mM ascorbate, 40 nM Pc. Excitation light duration is 0.1 sec. (C) NaCl concentration shows no effect on the fast rate of P700^+^ reduction by Pc. Reaction mixture contains: 20 mM MES-Tris (pH=6.0); 0.05% α-DM and PSI containing 5 µg Chl. All samples contain 10 mM ascorbate, 40 nM Pc. Excitation achieved by a 10 μsec laser flash.

## Discussion

In PSI of oxygenic organisms, the binding site of the reduced electron carriers is mediated by both cytochrome c_6_ and Pc. Vascular plants developed specificity for Pc, while cyanobacteria and algae use both as viable electron donors. Three subunits of PSI PsaA, PsaB and PsaF are directly involved in Pc binding (*17*). It was reported that the PSI binding site involve both electrostatic and hydrophobic interactions (*8, 9*). The structure of Pc-PSI supercomplex reveals not only the electrostatic interactions reported before (*17*) but also hydrophobic interactions (Fig 1, B to E, Table 1). It was suggested that the observed kinetics can be explained by a rate-limiting conformational change that occurs in the Pc–PSI complex before the electron transfer takes places (*33, 34*). We found no significant structural alterations of PSI in response to Pc binding. This is in line with an alternative model, that explains the kinetic properties of the electron transfer between Pc and PSI, where the driving force of the intercomplex electron transfer is decreased to ΔG = -55 meV and tuned for an optimized turnover of PSI (*13*). The formation of the intermolecular PSI-Pc complex leads to an exclusion of water molecules as revealed via the structural data (Fig 1). Notably, this is in accordance with a low reorganization energy λ of about 418 meV, that was determined for the intermolecular electron transfer between Pc and PSI (*35*), indicating a rather hydrophobic environment at the contact side of both proteins. Thus, electron transfer between Pc and PSI is only coupled to a small reorganization of solvent molecules supporting fast interprotein electron transfer despite relatively low driving force. According to Marcus and Sutin (*36*), such an exclusion of solvent molecules would result in a decrease in standard entropy of reduction ΔS° of bound compared to soluble Pc mainly from changes in freedom of the rotation and liberation of the solvent molecules which surround the reaction partners. Taking advantage of the thermodynamic parameters such as ΔG (free energy change) and λ (the reorganization energy) and an empirical approximation for intraprotein electron transfer (*37*), an edge-to-edge distance R between Pc and PSI redox centers of 14.5 Å were estimated (*35*). This value is very close to the recent structural determination of this distance of 14.7 Å from Cu+ to P700 (*17*). Therefore, in conclusion, the thermodynamic and structural data are consistent, both supporting the exclusion of solvent molecules from the intermolecular electron transfer complex between PSI and Pc. Interestingly, the exclusion of solvent molecules from the intermolecular complex also has a consequence of the pH-dependent redox midpoint potential of Pc (*35*).

The importance of the hydrophobic, polar and electrostatic properties formed by the two α-helices *l’* and *j’* for binding of Pc to PSI was further investigated by site-directed mutagenesis of a Arg/Asp residue pair PsaA-Asp648/Arg647. PsaA-Asp648/Arg647 are close to the Trp pair (PsaA-Trp651 and PsaB-Trp627), where indole groups are arranged in a sandwich like structure above the P700 chlorophyll pair. An alignment of 114 eukaryotic and 36 cyanobacterial PsaA amino-acid sequences revealed a very high sequence similarity, especially for the part of the sequence representing the α-helix *l’* (fig S6). The residues PsaA-Arg647 and PsaA-Asp648 are conserved through all sequences, thus only in *Dinoflagellate* the PsaA-Asp648 is substituted by Asparagine. The single flash absorption measurements revealed diminished second order rate constants as compared to wild type. The lowest rate was measured for PsaA-Asp648Arg (6.539 e^+07^ k_2_ / M^-1^ s^-1^). On the other hand, K_D_ values could not be determined for the electron transfer between Pc and altered PSIs as no or only a minor increase of the A(1) amplitude with increasing Pc concentration was observed. Thus, indicating that the K_D_ value is substantially increased due to destabilized complex formation between Pc and the altered PSIs. A similar effect was observed for the PsaB-Trp627Phe mutant PSI, where the secondary electron transfer rate was also only slightly diminished, whereas the affinity was strongly impaired. Thus, stable complex formation between PS and Pc is not required for rather productive electron transfer, as revealed by less severe impact on second order electron transfer constants. However, an exact positioning of electrostatic and polar properties of PsaA-Arg647/Asp648 are required for stable binding of Pc supporting the importance for a precise polar, electrostatic and hydrophobic landscape required for stable complex formation between PSI and Pc. This is also supported by the fact, that high salt concentrations did not impact the formation of the first-order electron transfer complex, while the second order electron transfer constant was diminished (table S4). This clearly underpins the significance of hydrophobic interactions for stable complex formation between PSI and Pc. The importance of a stable intermolecular electron transfer complex between PSI and Pc is signified by a severe growth phenotype under high light of mutant strains PsaA-D648R, PsaA-R647D and PsaA-R647D/D648R (fig S7). While the polar, electrostatic and hydrophobic landscape is crucial for tight and stable association between PSI and reduced Pc, it appears to be the weakening of the hydrophobic interaction caused by oxidation of Pc (movie S2), that allows water molecules to accommodate the space between Pc and PSI and that drives Pc dissociation from PSI. This in turn likely explains the six times larger dissociation constant of oxidized Pc as compared to reduced Pc (*13*), responsible for the 50-60 mV higher midpoint redox potential of Pc bound to PSI.

In summary, the electron transfer between Pc and PSI can be described by the following steps, (i) binding of Pc to PSI facilitated via electrostatic interactions between Pc and PSAF, (ii) stabilization of the reduced Pc-PSI complex due to the hydrophobic as well as polar and electrostatic interface provided by the PsaA and PsaB association domains, (iii) electron transfer from Pc to P700^+^ within the intermolecular complex, (iv) dissociation of bound oxidized Pc driven via its conformational change, that destabilizes its hydrophobic interaction with PsaA and PsaB interfaces additionally supported through electrostatic pulling towards PSAF.

In conclusion, our data provide mechanistic insights at molecular resolution revealing how the light driven electron transfer between Pc and PSI was optimized for fast turnover.

## Acknowledgements

Dr Yael Levi-Kalisman is gratefully acknowledged and thanked for vitrifying the samples. We thank Dr. Alexey Amunts for data collection in the SciLifeLab Cryo-EM facility, Solna, Sweden. We would like to thank Kevin Redding from the Department of Chemistry and Biochemistry of the Arizona state University for providing the Δ*psaA* strain as well as the plasmid used for site directed mutagenesis and transformation. Molecular graphics and analyses were performed with UCSF Chimera, developed by the Resource for Biocomputing, Visualization, and Informatics at the University of California, San Francisco, with support from NIH P41-GM103311. Funding: This work was supported by The Israel Science Foundation (Grant No. 569/17), and by German-Israeli Foundation for Scientific Research and Development (GIF) to N.N and M.H., Grant no. G-1483-207/2018. M.H. acknowledges support from German Science Foundation (DFG, Hi 739/13-2). N.B-T acknowledges the support of NSF-BSF grant 2019658 and Abraham E. Kazan Chair in Structural Biology, Tel Aviv University.

## Author Contributions

I.C., M.F., S.K., A.B.S., D.K., and N.N. performed the research. I.C., F.D., M.H. and N.N. analyzed the data. I.C., M.F., M.H. and N.N. wrote the manuscript. All the authors discussed, commented on and approved the final manuscript.

## Data and materials availability

The atomic coordinates have been deposited in the Protein Data Bank, with accession code 6ZOO. The cryo-EM maps have been deposited in the Electron Microscopy Data Bank, with accession codes EMD-11326. All other data are presented in the main text or supplementary materials.

## Supplementary Material

Materials and Methods

Figures S1-S8

Tables S1-S4

Movies S1-S2

References (*38-56*)

## References and Notes

1. N. Nelson, C. Yocum, Structure and Function of Photosystems I and II. Annu. Rev. Plant Biol. 57, 521–565 (2006). doi: https://doi.org/10.1146/annurev.arplant.57.032905.105350

2. H. Kirchhoff, Structural changes of the thylakoid membrane network induced by high light stress in plant chloroplasts. Phil. Trans. R. Soc. B 369: 20130225. (2014). http://dx.doi.org/10.1098/rstb.2013.0225

3. X. Liu, Y. Zhou, J. Xiao, F. Bao, Effects of Chilling on the Structure, Function and Development of Chloroplasts. Front. Plant Sci. 9:1715. (2018). doi: 10.3389/fpls.2018.01715

4. V. K. Dalal, B. C. Tripathy, Water-stress induced downsizing of light-harvesting antenna complex protects developing rice seedlings from photo-oxidative damage. Scientific reports. 8:5955 (2018). doi:10.1038/s41598-017-14419-4

5. N. Nelson, W. Junge, Structure and Energy Transfer in Photosystems of Oxygenic Photosynthesis. Annu. Rev. Biochem. 84, 659–683 (2015). doi: https://doi.org/10.1146/annurev-biochem-092914-041942

6. R. Höhner, M. Pribil, M. Herbstová, L. S. Lopez, H. H. Kunz, M. Li, M. Wood, V. Svoboda, S. Puthiyaveetil, D. Leister, H. Kirchhoff, Plastocyanin is the long-range electron carrier between photosystem II and photosystem I in plants. Proc Natl Acad Sci U S A. 117(26):15354–15362 (2020). doi: 10.1073/pnas.2005832117

7. H. Kirchhoff, M. A. Schöttler, J. Maurer, E. Weis, Plastocyanin redox kinetics in spinach chloroplasts: evidence for disequilibrium in the high potential chain. Biochim Biophys Acta. 1659(1):63–72 (2004.) doi: 10.1016/j.bbabio.2004.08.004

8. W. Haehnel, T. Jansen, K. Gause, R. B. Klösgen, B. Stahl, D. Michl, B. Huvermann, M. Karas, R. G. Herrmann, Electron transfer from plastocyanin to photosystem I. EMBO J. 13: 1028–1038 (1994). doi: https://doi.org/10.1002/j.1460-2075.1994.tb06351.x

9. M. Hippler, J. Reichert, M. Sutter, E. Zak, L. Altschmied, U. Schröer, R. G. Herrmann, W. Haehnel, The plastocyanin binding domain of photosystem I. EMBO J. 15:6374–84 (1996). doi: https://doi.org/10.1002/j.1460-2075.1996.tb01028.x

10. S. Kuhlgert, F. Drepper, C. Fufezan, F. Sommer, M. Hippler, Residues PsaB Asp612 and PsaB Glu613 of photosystem I confer pH-dependent binding of plastocyanin and cytochrome c(6). Biochemistry. 51, 7297–303 (2012). doi: 10.1021/bi300898j

11. U. Schreiber, Redox changes of ferredoxin, P700, and plastocyanin measured simultaneously in intact leaves. Photosynth Res. 134: 343–360 (2017). doi: 10.1007/s11120-017-0394-7

12. A. Busch, M. Hippler, The structure and function of eukaryotic photosystem I. Biochimica Biophysica Acta. 1807, 864–877 (2011). doi: https://doi.org/10.1016/j.bbabio.2010.09.009

13. F. Drepper, M. Hippler, W. Nitschke, W. Haehnel, Binding dynamics and electron transfer between plastocyanin and photosystem I. Biochemistry. 35, 1282–1295 (1996). doi: https://doi.org/10.1021/bi951471e

14. G. Shimakawa, C. Miyake, Oxidation of P700 ensures robust photosynthesis. Frontiers in plant science. 9, 1617 (2018).

15. M. G. Müller, J. Niklas, W. Lubitz, A. R. Holzwarth, Ultrafast transient absorption studies on Photosystem I reaction centers from Chlamydomonas reinhardtii. 1. A new interpretation of the energy trapping and early electron transfer steps in Photosystem I. Biophys J. 85(6), 3899–3922 (2003). doi:10.1016/S0006-3495(03)74804-8

16. L. Tian L, P. Xu, V. U. Chukhutsina, A. R. Holzwarth, R. Croce, Zeaxanthin-dependent nonphotochemical quenching does not occur in photosystem I in the higher plant Arabidopsis thaliana. Proc Natl Acad Sci U S A. 114(18), 4828–4832 (2017). doi: 10.1073/pnas.1621051114

17. I. Caspy, A. Borovikova-Sheinker, D. Klaiman, Y. Shkolnisky, N. Nelson, The structure of a triple complex of plant photosystem I with ferredoxin and plastocyanin. Nature Plants. 6, 1300–1305 (2020). doi: https://doi.org/10.1038/s41477-020-00779-9

18. F. Sommer, F. Drepper, M. Hippler, The luminal helix l of PsaB is essential for recognition of plastocyanin or cytochrome c6 and fast electron transfer to photosystem I in Chlamydomonas reinhardtii. J Biol Chem. 277, 6573–6581 (2002).

19. F. Sommer, F. Drepper, W. Haehnel, M. Hippler, The hydrophobic recognition site formed by residues PsaA-Trp651 and PsaB-Trp627 of photosystem I in Chlamydomonas reinhardtii confers distinct selectivity for binding of plastocyanin and cytochrome c6. J Biol Chem. 279, 20009–20017 (2004).

20. M. Hippler, F. Drepper, W. Haehnel, J. D. Rochaix, The N-terminal domain of PsaF: precise recognition site for binding and fast electron transfer from cytochrome c6 and plastocyanin to photosystem I of Chlamydomonas reinhardtii. Proc Natl Acad Sci U S A. 95(13), 7339–44 (1998). doi: 10.1073/pnas.95.13.7339. doi: https://doi.org/10.1073/pnas.95.13.7339

21. K. Sigfridsson, S. Young, Hansson, O, Electron Transfer Between Spinach Plastocyanin Mutants and Photosystem 1. European Journal of Biochemistry. 245, 805–812 (1997). doi:10.1111/j.1432-1033.1997.00805.x

22. K. Sigfridsson, S. Young, S. O. Hansson, Structural dynamics in the plastocyanin-photosystem 1 electron-transfer complex as revealed by mutant studies. Biochemistry, 35, 1249–1257 (1996). doi: https://doi.org/10.1021/bi9520141

23. H. Jansson, M. Okvist, F. Jacobson, M. Ejdebäck, O. Hansso, L .Sjölin, The crystal structure of the spinach plastocyanin double mutant G8D/L12E gives insight into its low reactivity towards photosystem 1 and cytochrome f. Biochim Biophys Acta. 1607(2-3), 203–10 (2003). doi: 10.1016/j.bbabio.2003.09.011.

24. M. Ubbink, M. Ejdebäc, B. G. Karlsson, D. S. Bendall, The structure of the complex of plastocyanin and cytochrome f, determined by paramagnetic NMR and restrained rigid-body molecular dynamics. Structure. 6(3), 323–35 (1998). doi: 10.1016/s0969-2126(98)00035-5.

25. J. Illerhaus, L. Altschmied, J. Reichert, E. Zak, R. G. Herrmann,W. Haehnel, Dynamic interaction of plastocyanin with the cytochrome bf complex. J Biol Chem. 275, 17590–17595 (2000). doi: 10.1074/jbc.275.23.17590

26. R. Fraczkiewicz, W. Braun, Exact and Efficient Analytical Calculation of the Accessible Surface Areas and Their Gradients for Macromolecules. J. Comp. Chem. 19, 319 (1998). doi: https://doi.org/10.1002/(SICI)1096-987X(199802)19:3<319::AID-JCC6>3.0.CO;2-W

27. R. Fraczkiewicz, W. Braun, A New Efficient Algorithm for Calculating Solvent Accessible Surface Areas of Macromolecules. Presented at the Third Electronic Computational Chemistry Conference, Northern Illinois University, IL, November 1996

28. G. Finazzi, F. Sommer, M. Hippler, Release of oxidized plastocyanin from photosystem I limits electron transfer between photosystem I and cytochrome b6f complex in vivo. Proc Natl Acad Sci U S A. 102, 7031–7036 (2005). doi: https://doi.org/10.1073/pnas.0406288102

29. G. S. Kachalova, A. C. Shosheva, G. P. Bourenkov, A. A. Donchev, M. I. Dimitrov, H. D. Bartunik, Structural comparison of the poplar plastocyanin isoforms PCa and PCb sheds new light on the role of the copper site geometry in interactions with redox partners in oxygenic photosynthesis. J Inorg Biochem. 115, 174–181 (2012). doi: 10.1016/j.jinorgbio.2012.07.015

30. S. F. Altschul, W. Gish, W. Miller, E. W. Myers, D. J. Lipman, Basic local alignment search tool. J. Mol. Biol. 215, 403–410 (1990). doi: https://doi.org/10.1016/S0022-2836(05)80360-2

31. M. Hippler, F. Drepper, J. Farah, J. D. Rochaix. Fast electron transfer from cytochrome c6 and plastocyanin to photosystem I of Chlamydomonas reinhardtii requires PsaF. Biochemistry 36, 6343–6349 (1997). doi: 10.1021/bi970082c

32. T. Hiyama, B. Ke. Difference spectra and extinction coefficients of P700, Biochim. Biophys. Acta 267, 160–171 (1972). doi: https://doi.org/10.1016/0005-2728

33. H. Bottin, P. Mathis. Interaction of plastocyanin with photosystem I reaction center: A kinetic study by flash absorption spectroscopy. Biochemistry 24, 6453–6460 (1985). doi: https://doi.org/10.1021/bi00344a022

34. H. Bottin, P. Mathis. Turn-over of Electron-Donors in Photosystem-I - Double-Flash Experiments with Pea-Chloroplasts and Photosystem-I Particles. Biochim Biophys Acta 892, 91–98 (1987). doi: https://doi.org/10.1016/0005-2728(87)90251-9

35. M. Hippler, F. Drepper, W. Haehnel. [The Oxidizing Site of Photosystem I Modulates the Electron Transfer from Plastocyanin to P700+]. In From Light to Biosphere, P. Mathis, ed. (Amsterdam, N.L.: Kluwer Academic Publishers.), pp. 99-102 (1995).

36. R.A. Marcus, N. Suttin. Electron transfer in chemistry and biology. Biochim Biophys Acta 811, 265–322 (1985). doi: https://doi.org/10.1016/0304-4173(85)90014-X

37. C.C. Moser, J.M. Keske, K. Warncke, R.S. Farid, P.L. Dutton. Nature of biological electron transfer. Nature 355, 796–802 (1992). doi: 10.1038/355796a0

38. S. Q. Zheng, E. Palovcak, J. P. Armache, K. A. Verba, Y. Cheng, D. A. Agard, MotionCor2: anisotropic correction of beam-induced motion for improved cryo-electron microscopy. Nat. Methods 14, 331–332 (2017). doi: 10.1038/nmeth.4193

39. A. Rohou, N. Grigorieff, CTFFIND4: fast and accurate defocus estimation from electron micrographs. J. Struct. Biol. 192, 216–221 (2015). doi: 10.1016/j.jsb.2015.08.008

40. A. Eldar, B. Landa, Y. Shkolnisky, KLT picker: Particle picking using data-driven optimal templates. J Struct Biol. 210(2),107473 (2020). doi: 10.1016/j.jsb.2020.107473.

41. J. Zivanov, T. Nakane, B. O. Forsberg, D. Kimanius, W. J. H. Hagen, E. Lindahl, S. H. W. Scheres, New tools for automated high-resolution cryo-EM structure determination in RELION-3. eLife 7 (2018). doi: https://doi.org/10.7554/eLife.42166

42. P. D. Adams, P. V. Afonine, G. Bunkóczi, V. B. Chen, I. W. Davis, N. Echols, J. J. Headd, L.-W. Hung, G. J. Kapral, R. W. Grosse-Kunstleve, A. J. McCoy, N. W. Moriarty, R. Oeffner, R. J. Read, D. C. Richardson, J. S. Richardson, T. C. Terwilliger, P. H. Zwart, PHENIX: a comprehensive Python-based system for macromolecular structure solution. Acta Crystallogr. D. 66, 213–221 (2010). doi: https://doi.org/10.1107/S0907444909052925

43. P. Emsley, B. Lohkamp, W. G. Scott, K. Cowtan, Features and development of Coot. Acta Crystallogr. D. 66, 486–501 (2010). doi: 10.1107/S0907444910007493

44. V. B. Chen, W. B. Arendall III, J. J. Headd, D. A. Keedy, R. M. Immormino, G. J. Kapral, L. W. Murray, J. S. Richardson, D. C. Richardson, MolProbity: all-atom structure validation for macromolecular crystallography. Acta Crystallogr. D. 66, 12–21 (2010). doi: 10.1107/S0907444909042073

45. A. Kucukelbir, F. J. Sigworth, H. D. Tagare, Quantifying the Local Resolution of Cryo-EM Density Maps. Nat. Methods. 11, 63–65 (2014). doi: 10.1038/nmeth.2727

46. The PyMOL Molecular Graphics System, Version 1.2r3pre, Schrödinger, LLC.

47. E. F. Pettersen, T. D. Goddard, C. C. Huang, G. S. Couch, D. M. Greenblatt, E. C. Meng, T. E. Ferrin, UCSF chimera—a visualization system for exploratory research and analysis. J. Comput. Chem. 25, 1605–1612 (2004). doi: 10.1002/jcc.20084

48. Harris, E.H. (1989). The Chlamydomonas Sourcebook (San Diego: Academic Press).

49. N. H. Chua, P. Bennoun, Thylakoid membrane polypeptides of Chlamydomonas reinhardtii: wild-type and mutant strains deficient in photosystem II reaction center. Proc Natl Acad Sci U S A. 72, 2175–2179 (1975). doi: 10.1073/pnas.72.6.2175

50. Y. Takahashi, M. Goldschmidt-Clermont, S. Y. Soen, L. G. Franzen, J. D. Rochaix, Directed chloroplast transformation in Chlamydomonas reinhardtii: insertional inactivation of the psaC gene encoding the iron sulfur protein destabilizes photosystem I. Embo J. 10, 2033–2040 (1991). doi: https://doi.org/10.1002/j.1460-2075.1991.tb07733.x

51. S. Katoh, I. Shiratori, A. Takamiya, Purification and some properties of spinach plastocyanin. J Biochem. 51, 32-40. (1962a). doi: 10.1093/oxfordjournals.jbchem.a127497

52. L. Zheng, U. Baumann, J. L. Reymond, An efficient one-step site-directed and site-saturation mutagenesis protocol. Nucleic Acids Res. 32, e115 (2004). doi: 10.1093/nar/gnh110

53. U. K. Laemmli, Cleavage of structural proteins during the assembly of the head of bacteriophage T4. Nature. 227, 680–685 (1970). doi: https://doi.org/10.1038/227680a0

54. R. J. Porra, W. A. Thompson, P. E. Kriedemann, Determination of accurate extinction coefficients and simultaneous equations for assaying chlorophylls a and b extracted with four different solvents: verification of the concentration of chlorophyll standards by atomic absorption spectroscopy. Biochimica et Biophysica Acta – Bioenergetics. 975, 384–394 (1989). doi: https://doi.org/10.1016/S0005-2728(89)80347-0

55. R. C. Edgar, MUSCLE: multiple sequence alignment with high accuracy and high throughput. Nucleic acids research. 32, 1792–1797 (2004). doi: 10.1093/nar/gkh340

56. G. E. Crooks, G. Hon, J. M. Chandonia, S. E. Brenner, WebLogo: a sequence logo generator. Genome Res 14, 1188–1190 (2004). doi: 10.1101/gr.849004

