## Supplementary material for "Structure of plant PSI-plastocyanin complex reveals strong hydrophobic interactions": PSI_Pc_Supplementary_Materials_Final.pdf

### Materials and Methods

P700+ reduction:

In all the kinetics measurements described in this work about 50 large PSI crystals were collected using 0.7-1.0 mm Cryoloop (Hampton research) and released into a 0.5 ml Eppendorf tube containing 0.4 ml solution of 50 mM $K_2PO_4$, 50 mM Tris (pH8) and 20% PEG 400. Following centrifugation at 14,000 rpm for 1 min, the supernatant was carefully removed, and the crystals were solubilized in 60 µl of a solution containing 20 mM Tricine-Tris (pH8) and 0.05% αDM at a chlorophyll concentration of 1 to 3 mg/ml.

Cryo-EM data collection and processing:

For cryo-EM structural determination, about 50 large PSI crystals were collected using 0.7-1.0 mm Cryoloop (Hampton research) and released into a 0.5 ml Eppendorf tube containing 0.4 ml solution of 50 mM $K_2PO_4$, 50 mM Tris (pH8) and 20% PEG 400. Following centrifugation at 14,000 rpm for 1 min, the supernatant was carefully removed, and the crystals were solubilized in 60 µl of a solution containing 20 mM MES-Tris (pH7), 20 mM sodium ascorbate and 0.05% αDM at a chlorophyll concentration of 6 mg/ml. A concentrated pea plastocyanin was added into the PSI solution to give final concentration of PSI 2.8 mg Chl/ml and 5mg/ml plastocyanin. The resulting solution of PSI-Pc (3 µl) was applied on glow-discharged holey carbon grids (Cu Quantifoil R1.2/1.3), before using a Vitrobot FEI (3 s blot at 4°C and 100% humidity). The frozen grids were sent to Dr. Alexey Amunts at the SciLifeLab. Images were collected in the Karolinska Institute (KI) facility using a K3- 300 kV FEI Titan Krios electron microscope. The images were collected using a 300 kV FEI Titan Krios electron microscope, with an energy filter of 10 eV. A Gatan K3-Summit detector was used in fast acquisition mode at a magnification of

×130,000 (yielding a pixel size of 0.654 Å), with a total dose of 40.8 e $Å^{-2}$. EPU was used to collect a total of 5,310 images, which were dose-fractionated into 51 video frames, with defocus values ranging from 0.3 µm to 1.5 µm. The collected micrographs were motion-corrected and dose-weighted using MotionCor2 (*38*). The contrast transfer function parameters were estimated using CtfFind 4.1 (*39*). A total of 276,815 particles were picked using reference-free picking using the KLT picker (*40*). The picked particles were processed for reference-free 2D averaging, initial model building, 3D classification and refinement resulting in 45,376 particles for intermediate model building; All steps were performed using RELION v.3 (*41*). The intermediate model was subsequently used for template based autopicking in RELION, resulting in 359,977 picked particles. This set was used applied to 2D classification resulting in 141,959 particles, then followed by 3D classification resulting in six distinct classes in RELION. From these, four classes were selected, which contained a total of 104,127 particles. These particles were pooled together and processed for 3D homogeneous refinement and postprocessing using RELION. The reported resolution was based on a gold-standard refinement, applying the 0.143 criterion on the FSC between the reconstructed half-maps. Following data processing (see Methods section) the Pc-PSI supercomplex was modelled at 2.74 Å resolution with local resolution ranging from 2.5 to 4.5 (fig S1).

Model building:

To generate the Pc-PSI, the cryo-EM structure of the plant Pc-PSI–Fd model PDB 6YEZ was selected. This model was fitted onto the cryo-EM density map using phenix.dock_in_map in the PHENIX suite (*42*), and manually rebuilt using Coot (*43*). Stereochemical refinement was performed using phenix.real_space_refine in the PHENIX suite (*42*). The final model was validated using MolProbity (*44*). The refinement statistics are provided in table S1. Local resolution was determined using ResMap (*45*), and the figures were generated using PyMOL (*46*) and UCSF Chimera (*47*). Representative cryo-EM densities are shown in fig S8.

Strains and media:

Wild type and mutant strains were grown as described (*48*). Tris-acetat-phosphate (TAP) medium was used to grow cells for PSI isolation. The growth tests were performed on solidified (1.5 % agar) TAP medium for photoheterotrophic and high salt medium (HSM) for photoautotrophic growth.

Isolation of PSI particles:

Whole cells were used for Thylakoid membrane isolation according to established procedures (*49*). Afterwards the isolated membranes were diluted to a chlorophyll concentration of 0.8 mg/ml, solubilized and incubated on ice for 20 min in the presence of 0.9 % (w/v) dodecyl β-maltoside. The photosynthetic complexes were separated using a linear sucrose density gradient (*50*). The photosystem I was collected and concentrated by using ultra-filtration colums with a size exclusion of 100,000 MW.

Heterologous expression and isolation of Plastocyanin:

Plastocyanin was heterologously expressed as described  using the *E. coli* BL21 strain. The concentration of the isolated Pc was determined spectroscopically as described (*51*).

Growth tests:

The cell growth at different light conditions was determindet by spotting 20 µl of cells from log phase cultures, which were diluted to the same cell-number ($1*10^6$ cells / ml) onto agar plates. These plates were kept under 400 µE $m^{-2}$ $s^{-1}$ termed as high light, 40 µE $m^{-2}$ $s^{-1}$ as normal light and 5 µE $m^{-2}$ $s^{-1}$ as low light, respectively. The Genebag Anaerobiosis system provided by BioMérieux was used to establish anaerobic growth.

Generation of mutant strains:

The point mutations Asp647Arg, Arg648Asp and Asp647Arg/Arg648Asp in the photosystem I subunit PsaA were introduced by site directed mutagenesis (*52*). The transformation success was confirmed by sequencing as well as the expression and the expression level of the altered photosystem I by Western blot analysis.

SDS-PAGE and Western blot analysis:

SDS-PAGE (13 % T, 3.45 % C) was carried out as published (*53*). Proteins were transferred onto a nitro-cellulose membrane by semidry electro-blotting. The imuno detection was performed using anti-PsaD and anti-PsaF antibodies for PSI detection in a 1:1000 dilution as well as anti-D1 and anti-LhcbM6 antibodies of PSII detection in a 1:2000 and 1:1000 dilution, respectively.

Single flash absorbance spectroscopy:

The kinetics of flash-induced absorbance changes at 817 nm were measured as described before using a cuvette containing 200 µl of the sample with an optical path length of 10 mm. The chlorophyll content was determined according to the protocol published by Porra (*54*).

NADP-photoreduction measurements:

The light driven, PSI dependent reduction of $NADP^+$ was performed as described by Finazzi and co-workers (*38*). The sample was illuminated with saturating light supplied by a halogen lamp with a light intensity of about $10^4$ µE $m^{-2}$ $s^{-1}$.

Multiple Sequence Alignment (WebLogo):

150 annotated and reviewed sequences from various photosynthetic phyla where used to generate a multiple sequence alignment using MUSCLE (*55*) (sequences taken from UniProt). The WebLogo (*56*) was generated taking subsequence representing the PsaA α-helix l' from the previously generated alignment.

**Supplementary text**

PSI electron donors and acceptors

In addition to plastocyanin, P700 can accept electrons from a variety of non-native donors including organic redox compounds. At the opposite side of the complex, electrons can be withdrawn from PSI by non-native acceptors such as methylviologen. The organic redox compound Phenazine methosulfate (PMS) is highly efficient in donating and accepting electrons from PSI and thus is highly efficient in catalyzing cyclic photophosphorylation. PMS addition in the presence of ascorbic acid completely reduces the P700 population and cause significant changes in the light induced redox kinetics.

C. reinhardtii PSI mutant and growth

Notably, Asp648 (pea Asp655) is found at the beginning of the second PsaA surface. Both amino acids PsaA-Arg647 and Asp648 are found very close to Trp651 (pea Trp658) and are also present right at the PsaA-PsaB border. PsaA-Asp648 faces Pc Phe35 directly, and the salt bridge formed by Arg647 and Asp648 (fig S3) prevents them from drifting towards PsaA Trp658 or PsaB Asn633 (pea annotations), the latter found at the interface of the first Pc domain formed by Asp8-Leu12. The coding sequence for both residues in the *psaA* gene was altered to generate a scenario where both encoded residues are positively charged (PsaA-Asp648Arg), both are negatively charged, (PsaA-Arg647Asp) and where the charge is switched (PsaA-Arg647Asp/Asp648Arg). The *psaA* plasmid coding for the altered containing the *addA* gene (encoding for aminoglycoside adenyl transferase and conferring resistance to spectinomycin or streptomycin) was transformed into to a PsaA-deficient C. reinhartii mutant strain. The generated strains were selected on spectinomycin-containing medium and the amounts of expressed PSI (using anti-PsaD antibodies) were investigated.

Growing the mutant strains on solidified media under different light conditions uncovered a severe light sensitivity under photoautotrophic conditions (HSM – Minimal Media) for all light intensities and under photoheterotrophic conditions (TAP – Acetate containing media) at high light intensities (fig S6). In contrast, cells growing under photoheterotrophic conditions at normal or low light showed only slightly impaired growth. This phenotype is independent from the copper availability in the media. Under anoxic conditions, the growth phenotype was rescued for all strains, implying ROS formation caused the pronounced light sensitivity (fig S6), as reported for other C. reinhardtii mutants with impaired electron transfer between PSI and Pc.

<u>Reduction kinetics of light induced P700+</u>

The light induced absorption change at 705 nm measured by JTS that was calibrated by using plant PSI crystals containing 156 chlorophyll molecules per P700. The standard variation of Chl/P700 ratio from five independent experiments is ± 2%, caused mainly due to variation in chlorophyll determination and not the extent of P700 signal.

The data revealed that the reduction of light induced oxidized P700+ is biphasic with a fast kinetics of about 100 milliseconds and a slow one of several minutes. Addition of ferredoxin eliminated the fast kinetics. Ascorbate failed to reduce all the population of P700+ and the idle P700 disappeared in the presence of PMS and thus the signal represents the entire P700 population. The presence of PMS eliminated the fast 10 ms back kinetics that was attributed to FB. Only partial elimination of the back reaction was observed in the presence of Pc.

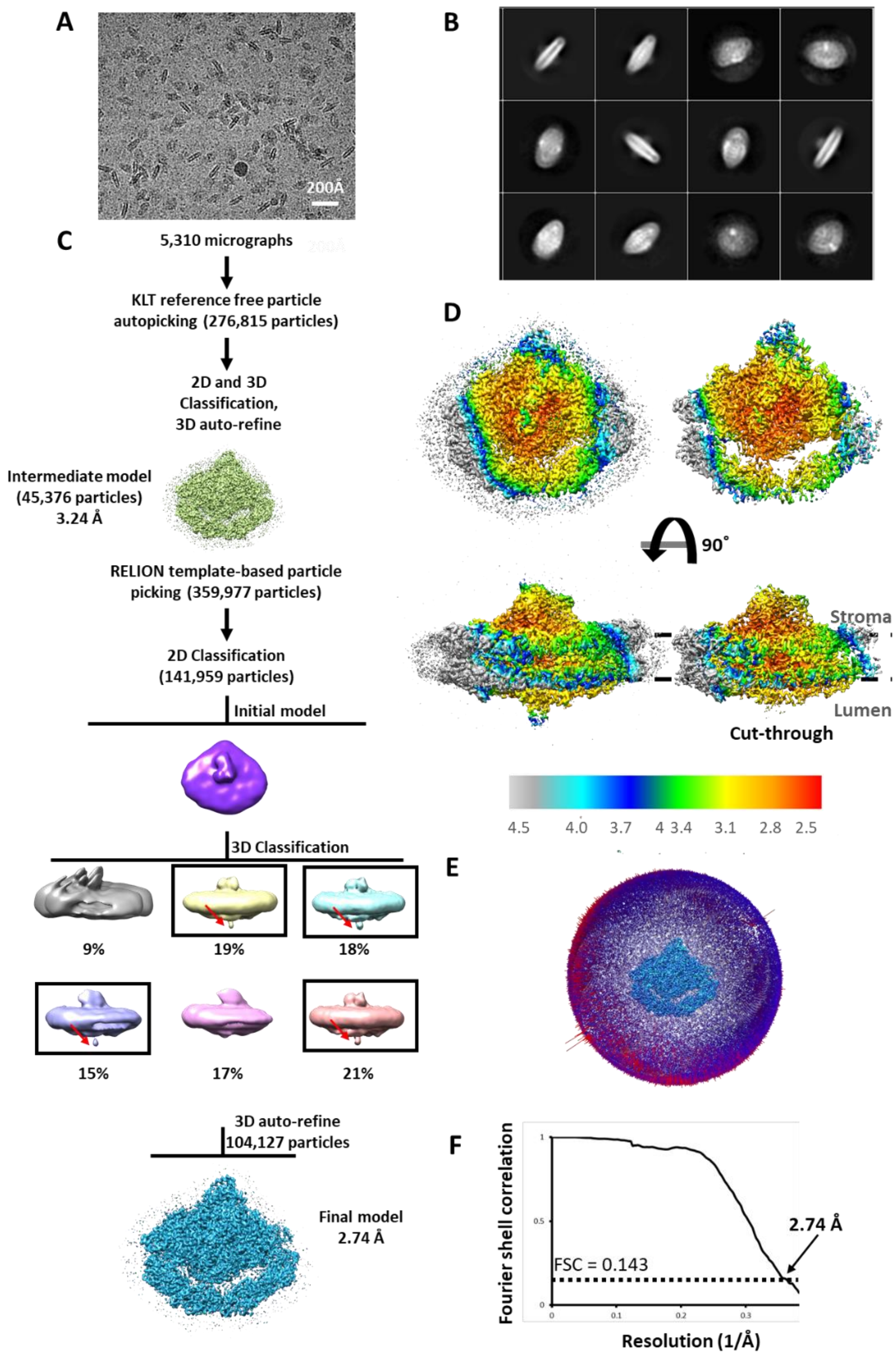


**Fig. S1.**

Cryo-EM data collection and processing of PSI-Pc. (A) A zoomed-in view on a PSI-Pc micrograph. Scale bar size is 200 Å. (B) Representing class averages after 2D classification. (C) Cryo-EM data processing workflow. Red arrows in the 3D classes indicate bound Pc. (D) Local resolution of the complete PSI-Pc (left) and map cut-through (right). (E) Angular distribution of PSI-Pc. (F) Fourier shell correlation (FSC) RELION postprocessing result marks a 2.74 Å resolution based on the 0.143 gold-standard.

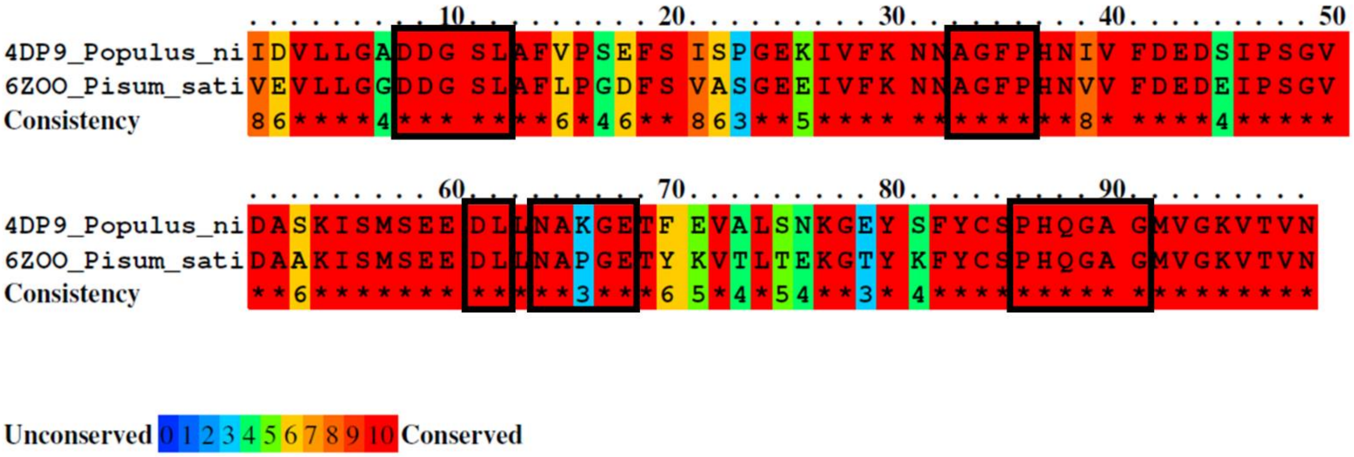


**Fig. S2.**

Sequence alignment of Pc from *Populus nigra* (PDB 4DP9) and *Pisum sativum* (PDB 6ZOO) using ClustalW. Rectangles mark the amino acids at the PsaA-PsaB interface as identified by SASA analysis.

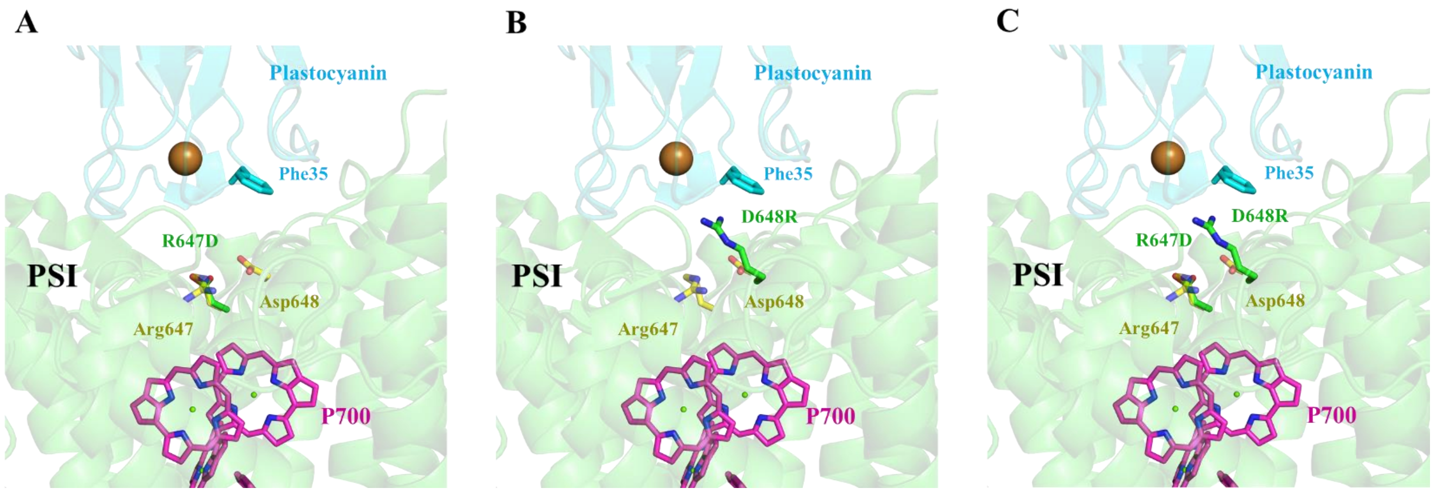


**Fig. S3.**

PsaA mutations impair Pc association with PSI. WT amino acids are colored yellow, mutations in green, Pc in cyan, P700 in magenta. Pc copper is shown as a brown sphere. (A) Arg647Asp (R647D) mutant, creating two negative charges near Pc. (B) Asp648Arg (D648R) mutant, forming two positive charges close to Pc with an additional interference with Pc Phe35. (C) Arg647Asp- Asp648Arg (R647D/D648R) mutant shows Arg648 clashes with Pc Phe35.

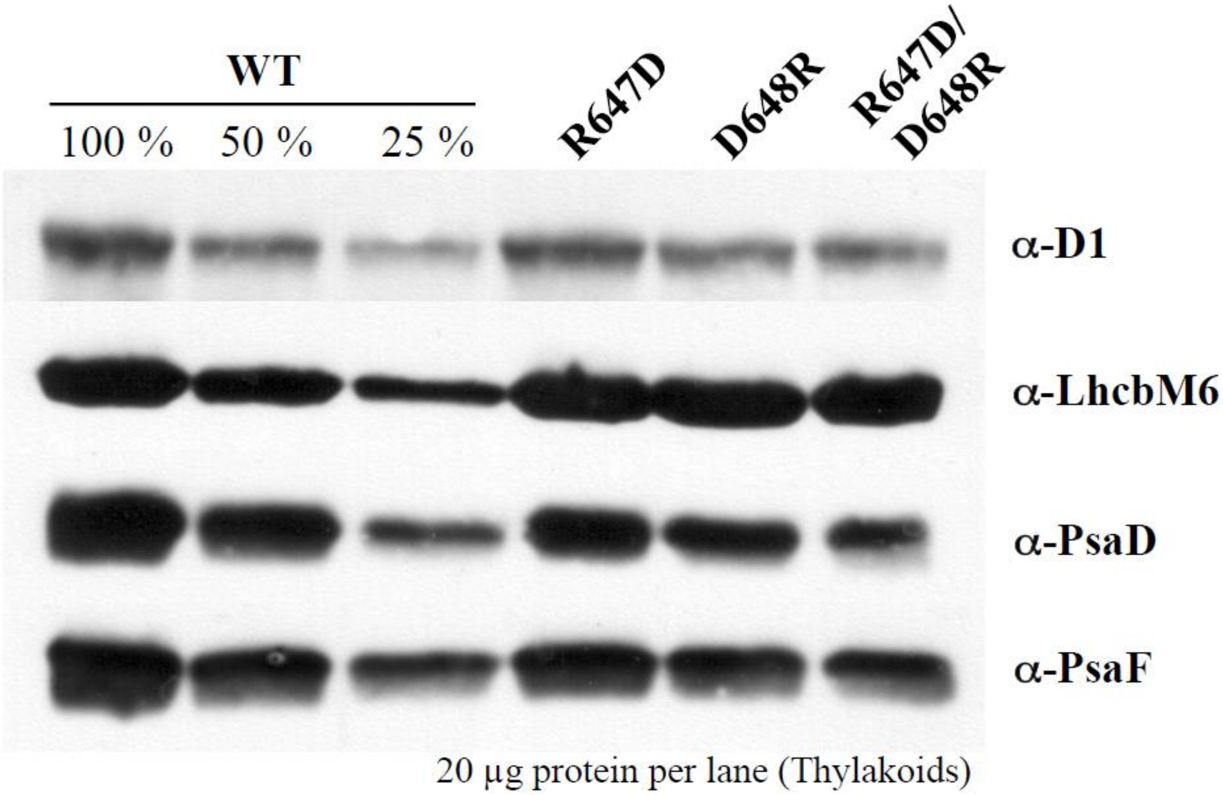


**Fig. S4.**

Western blot analysis of isolated Thylakoid membranes loaded on equal protein (20 μg protein / lane) and a dilution series for WT was made. The mutants are showing only slightly reduced amounts of D1 and LhcbM6 representing PSII, whereas the amounts of PsaD and PsaF representing PSI are only 25% to 50% of wild type level.

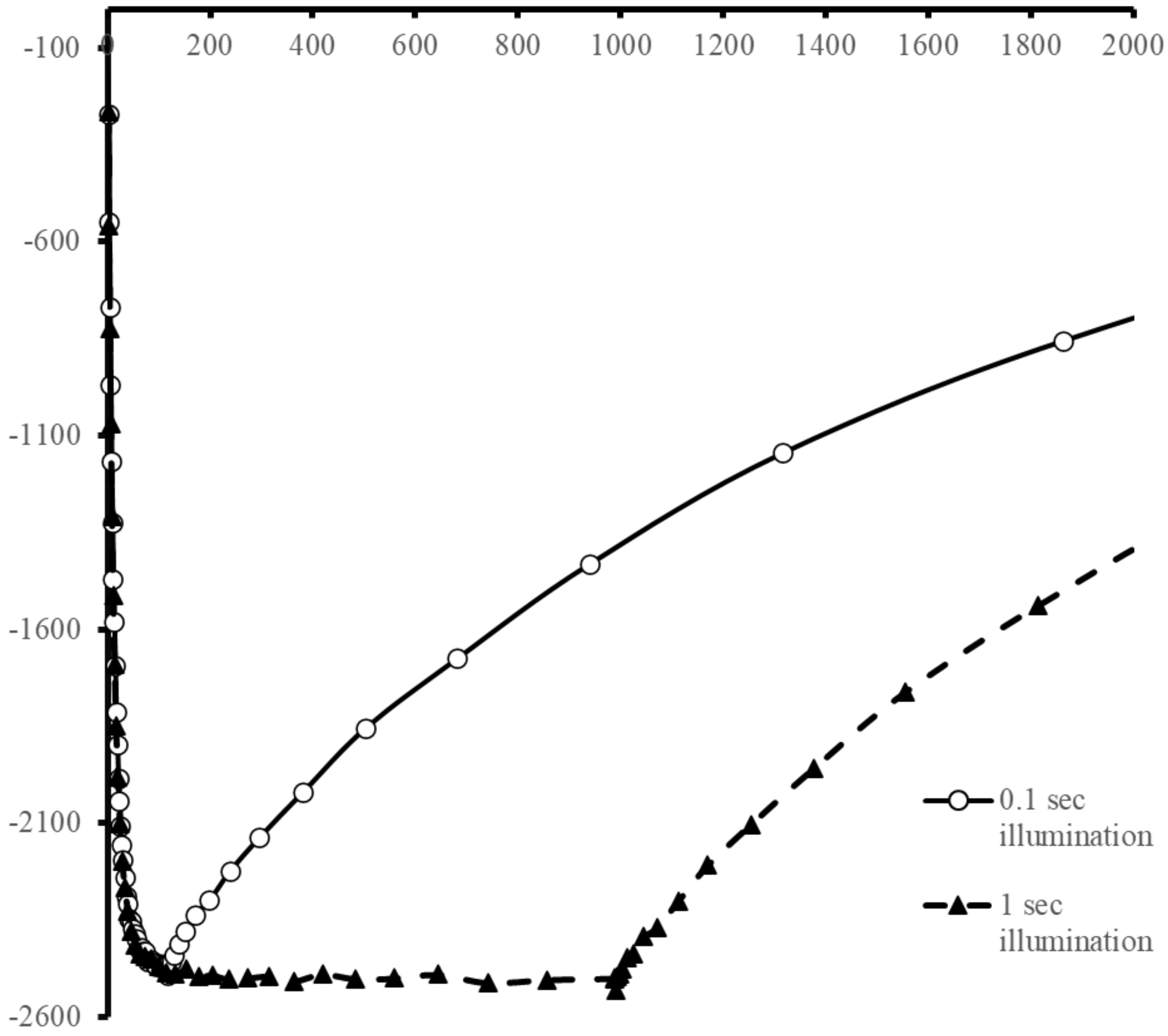


**Fig. S5.**
P700 OD remains stable upon illumination duration increase.

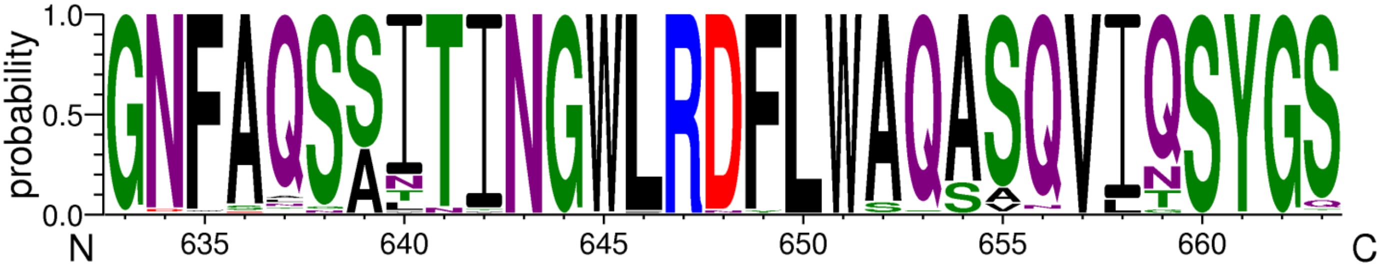


**Fig. S6.**

WebLogo generated for the PsaA α-helix l' from an alignment of 150 annotated and reviewed sequences from different photosynthetic phyla available on UniProt. The residues R647 and D648 are conserved through all sequences except for the ones from *Dinoflagellata* where the aspartic acid at position PsaA-648 is replaced by asparagine.

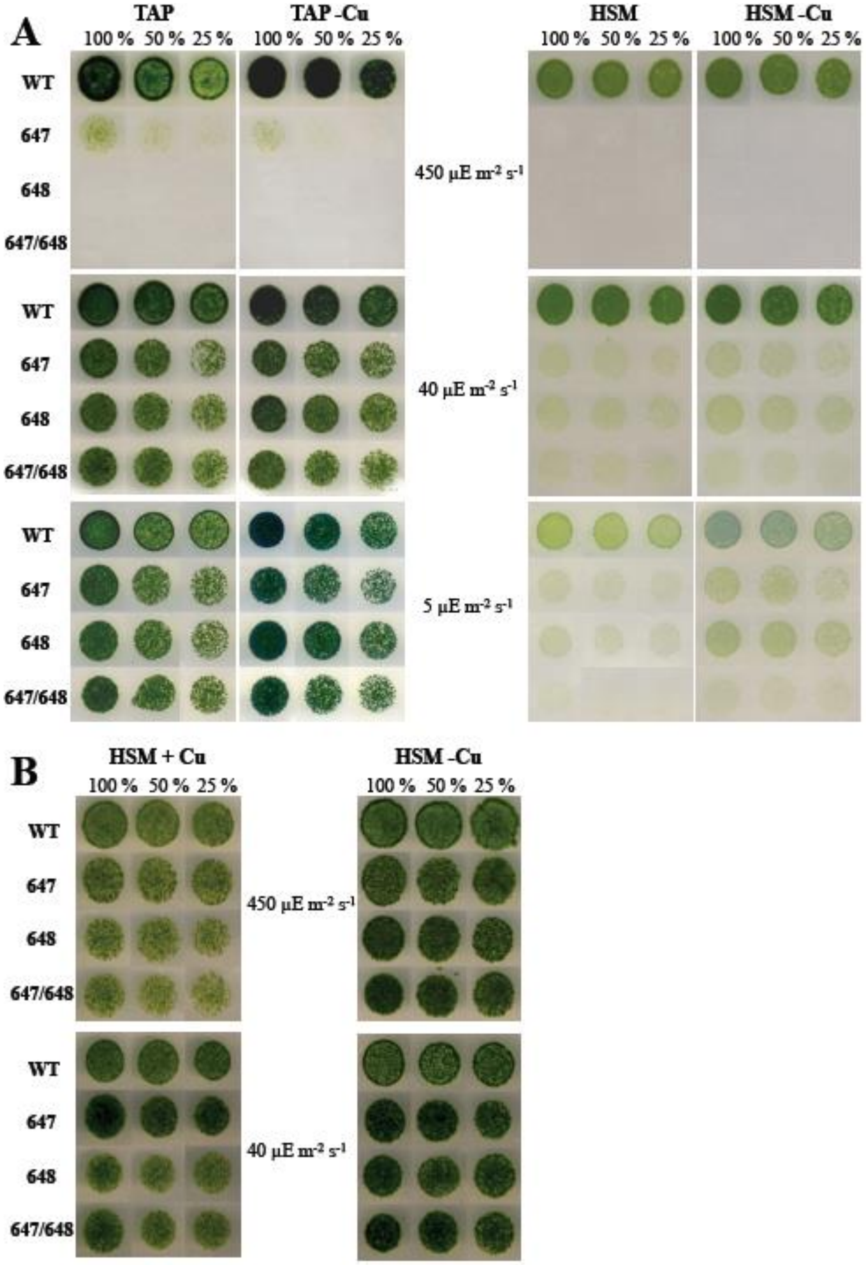


**Fig. S7.**

The wild type and mutant strains R647D, D648R and R647D/D648R were grown under different light intensities under photoheterotrophic (TAP media) and photoautotrophic (HS media) conditions. All media where also prepared without Copper (-Cu) to induce expression of cyt c6. (A) All mutant strains are showing a severe light sensitive growth phenotype under photoautotrophic conditions independent from the light intensity and at high light (450 µE m-2 s-1) under photoheterotrophic conditions. (B) Growing the mutant strains under anaerobic photoautotrophic conditions rescues the severe light sensitive growth phenotype for all used light intensities, independently from the electron donor pc (copper sufficient media) and cyt c6 (copper deficient media).

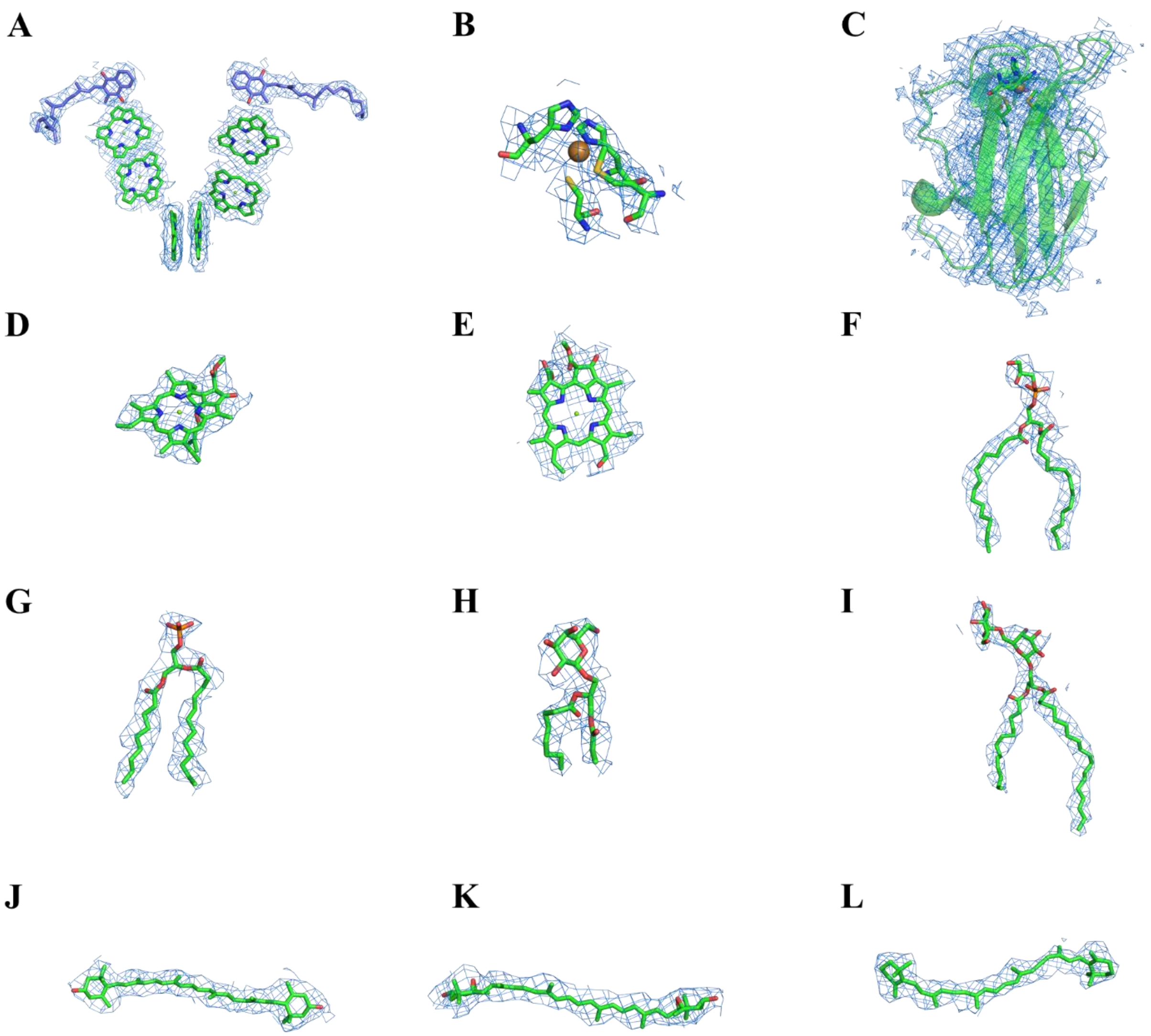


**Fig. S8.**

Cryo-EM densities of Pc and selected PSI ligands. (A) Cryo-EM densities of the electron transport chain main cofactors. Chlorophylls are colored green, quinones in marine. (B) Pc copper ion and coordinating amino acids. (C) Plastocyanin complete cryo-EM densities. Copper binding is shown as sticks. (D) Representing chlorophyll a (CLA), PsaL 1501. (E) Representing chlorophyll b (CHL), Lhca3 611. (F) Phosphatidylglycerol (PG), PsaA 5002. (G) Phosphatidic acid (PA), Lhca2 807. (H) Monogalactosyldiacylglycerol (LMG), PsaF 5001. (I) Digalactosyldiacylglycerol (DGD), PsaB 5005. (J) Lutein (LUT), Lhca1 and 501. (K) Violaxanthin (XAT), Lhca2 502. (L) β-carotene (BCR), PsaB 4010.

**Table S1.**
Cryo-EM data collection, refinement and validation statistics.

| | |
|---|---|
| **Data collection and processing** | |
| Magnification | 130,000 |
| Voltage (kV) | 300 |
| Electron exposure | 40.8 |
| Defocus range (μm) | 0.3-1.5 |
| Pixel size (Å) | 0.654 |
| Symmetry imposed | C1 |
| Micrographs | 5,310 |
| Initial particles images (no.) | 359,977 |
| Final particles images (no.) | 104,127 |
| Map resolution (Å) | 2.74 |
| FSC threshold | 0.143 |
| Map-sharpening B-factor ($Å^2$) | -38.8 |
| **Refinement** | |
| Initial model used (PDB code) | 6YEZ |
| Model resolution | 2.74 |
| FSC threshold | 0.143 |
| Model resolution range (Å) | 2.5-4.5 |
| Model composition | |
| Non-hydrogen atoms | 38,506 |
| Protein residues | 3339 |
| Ligands | 245 |
| B-factors ($Å^2$) | |
| Protein | 49.52 |
| Ligand | 49.92 |
| R.m.s deviations (PHENIX) | |
| Bond length (Å) | 0.013 |
| Bond angles (º) | 2.017 |
| Validation | |
| MolProbity Score | 2.29 |

| | |
|---|---|
| Clashscore | 24.21 |
| Poor rotamers (%) | 0.04 |
| Ramachandran plot | |
| Favored (%) | 93.67 |
| Allowed (%) | 6.03 |
| Disallowed (%) | 0.30 |

**Table S2.**

SASA of the in-silico model of PSI bound to oxidized Pc.
Total SASA for each residue in Å$^2$; Ratio – exposed SASA; Change – difference in SASA for each residue in complex or unbound. Charged or polar amino acids are colored blue, hydrophobic amino acids colored yellow.

| PsaA-PsaB total SASA | PsaA-PsaB in-complex total SASA | Pc total SASA | Pc in-complex total SASA |
|---|---|---|---|
| 69611 | 69149 | 5024 | 4539 |

| Residue | Chain | Number | PSI-Pc complex | | PSI-Pc unbound | | Change% |
|---|---|---|---|---|---|---|---|
| | | | Total SASA | Ratio% | Total SASA | Ratio% | |
| LEU | Pc | 12 | 49 | 27 | 60 | 35 | -8 |
| GLY | | 34 | 34 | 39 | 40 | 46 | -7 |
| PRO | | 36 | 9 | 8 | 62 | 58 | -50 |
| GLU | | 60 | 68 | 43 | 146 | 86 | -42 |
| ASP | | 61 | 64 | 53 | 70 | 54 | -1 |
| LEU | | 62 | 23 | 3 | 75 | 39 | -36 |
| ASN | | 64 | 18 | 7 | 89 | 64 | -57 |
| ALA | | 65 | 6 | 8 | 38 | 55 | -47 |
| PRO | | 66 | 79 | 62 | 94 | 77 | -14 |
| GLY | | 67 | 37 | 19 | 61 | 35 | -16 |
| GLU | | 68 | 26 | 14 | 27 | 16 | -2 |
| SER | | 85 | 25 | 3 | 79 | 37 | -34 |
| PRO | | 86 | 7 | 4 | 31 | 19 | -15 |
| HIS | | 87 | 57 | 65 | 86 | 99 | -34 |
| GLY | | 89 | 61 | 26 | 73 | 44 | -18 |
| ALA | | 90 | 562 | 27 | 1032 | 35 | -8 |
| | | | 49 | | 60 | | |
| THR | PsaA | 635 | 82 | 57 | 110 | 84 | -27 |
| ILE | | 637 | 24 | 6 | 55 | 7 | -1 |
| THR | | 638 | 19 | 14 | 27 | 16 | -3 |
| GLY | | 639 | 46 | 53 | 80 | 92 | -39 |
| TRP | | 658 | 36 | 14 | 66 | 27 | -13 |
| ALA | | 659 | 21 | 33 | 40 | 51 | -19 |
| SER | | 662 | 2 | 3 | 22 | 29 | -26 |
| GLN | | 663 | 6 | 1 | 46 | 28 | -27 |
| GLN | | 666 | 37 | 19 | 83 | 51 | -32 |
| GLY | | 669 | 43 | 49 | 43 | 49 | 0 |
| SER | | 670 | 21 | 18 | 53 | 59 | -41 |
| LEU | | 672 | 20 | 14 | 28 | 19 | -5 |
| ARG | | 753 | 5 | 2 | 19 | 10 | -7 |
| ALA | | 756 | 26 | 4 | 49 | 27 | -23 |
| VAL | | 757 | 32 | 7 | 52 | 23 | -16 |
| ASN | | 605 | 40 | 33 | 58 | 48 | -15 |

| | | | | | | | |
|---|---|---|---|---|---|---|---|
| **GLN** | **PsaB** | 608 | 33 | 20 | 45 | 29 | -9 |
| **TRP** | | 625 | 22 | 7 | 48 | 19 | -12 |
| **LEU** | | 626 | 45 | 27 | 86 | 55 | -28 |
| **SER** | | 629 | 33 | 42 | 38 | 49 | -7 |
| | | | 596 | | 1050 | | |

**Table S3.**
Determined second order rate constants ($k_2$ / $M^{-1}$ $s^{-1}$) and dissociation constants ($K_D$) for Pc and PSI at 1 mM $MgCl_2$. The PSI particles were isolated from wild type and altered PSI from mutant strain.

| | PC | | |
|---|---|---|---|
| | $k_2$ / $M^{-1}$ $s^{-1}$ | $K_D$ [μM] | *f* |
| Wild type | 3.792 e+07 | 88 | 0,50 |
| R647D | 1.504 e+07 | ------- | ------- |
| D648R | 6.539 e+06 | ------- | ------- |
| R647D/D648R | 1.798 e+07 | ------- | ------- |

**Table S4.**
NADP-photoreduction-rates in µM/s for wild type and altered photosystem particles in the presence of 1.25 µM and 80 µM recombinant wild type Pc, respectively. Three independent biological replicates were performed.

| PSI | PC [1.25 µM] | PC [80 µM] |
|---|---|---|
| Wild type | 0.8269 +/- 0. 0317 | 4.8553 +/- 0. 1739 |
| R647D | 0.2803 +/- 0. 0474 | 3.1972 +/- 0. 0490 |
| D648R | 0.3258 +/- 0. 0134 | 2.1329 +/- 0. 1774 |
| R647D/D648R | 0.1635+/- 0. 0188 | 3.8907 +/- 0. 2069 |

**Movie S1.**

Plastocyanin association to the hydrophobic interface formed by PsaA-PsaB as well as a small number of electrostatic interactions with PsaF. Pc, PsaA, PsaB and PsaF are shown as coulombic surfaces.

**Movie S2.**

In-silico model of proposed Pc dissociation from PSI. Pc acidic eastern patches Asp42-Glu45 and Glu59-Glu60 undergo a conformational change, bringing them closer to PsaF Lysines. This in turn causes Pc to tilt and drift towards PsaF and begin its dissociation from PSI, allowing water molecules to enter the solvent inaccessible surface formed by Pc binding to PsaA-PsaB. Pc is colored bright blue, copper as sphere and coordinates amino acids shown as sticks, PsaF in cyan, ETC chlorophylls colored tan. PSI subunits are shown as transparent cartoons, PsaA in green and PsaB in blue.
